# Modeling Hierarchical Brain Dynamics Outperforms Hormonal Biomarkers in Predicting Menstrual Cycle Phases

**DOI:** 10.64898/2025.12.01.691604

**Authors:** Elvira del Agua, Daniela Avila-Varela, Esmeralda Hidalgo-Lopez, Paulina Clara Dagnino, Marian Martínez-Marín, Irene Acero-Pousa, Yonatan Sanz Perl, Morten Kringelbach, Gustavo Deco, Belinda Pletzer, Anira Escrichs

**Author notes:** These authors contributed equally (Elvira del Agua, Daniela Avila-Varela). These authors share senior authorship (Belinda Pletzer, Anira Escrichs).

## Abstract

Hormonal fluctuations across the menstrual cycle influence large-scale brain dynamics, yet the underlying neurobiological mechanisms remain poorly understood. In this study, 60 naturally cycling women were scanned using resting-state fMRI during the early follicular, pre-ovulatory, and mid-luteal phases. We then applied a thermodynamics-inspired framework to explore the functional hierarchical organization of whole-brain dynamics across these phases. First, we found that brain dynamics are significantly modulated by estradiol, progesterone, and age across multiple resting-state networks. Second, to elucidate underlying mechanisms, we estimated generative effective connectivity (GEC) matrices using whole-brain models and trained support vector machine classifiers to predict menstrual phases. These model-based biomarkers outperformed traditional functional connectivity and hormone measures in classifying menstrual cycle phases. These findings reveal that menstrual cycle-related changes modulate the hierarchical reorganization of brain dynamics, highlighting the potential of model-based approaches to advance women’s brain health research.

## Introduction

The menstrual cycle, a fundamental physiological process in women of reproductive age, is orchestrated by dynamic fluctuations in ovarian hormones, primarily estradiol and progesterone [1]. Typically spanning 21 to 35 days [2], the cycle is primarily characterized by two distinct phases: the follicular phase, marked by rising levels of estradiol, and the luteal phase, distinguished by the dominance of progesterone and a secondary increase in estradiol. The transition between these two phases is a singular event: ovulation, which is triggered by a surge in luteinizing hormone (LH) that immediately follows the primary peak in estradiol. [3].

As an endocrine organ, the brain is highly sensitive to hormonal fluctuations throughout the menstrual cycle. Neuroimaging studies have consistently demonstrated menstrual cycle-related changes in brain connectivity patterns [4–11]. During the early follicular phase, brain dynamics are typically characterized by reduced information transmission, lower dynamical complexity, and greater temporal stability relative to the pre-ovulatory and mid-luteal phases [5, 7, 9]. In contrast, the ovulatory phase is associated with heightened dynamical complexity, increased brain flexibility, and reduced stability of brain connectivity (i.e., greater temporal variability) [7–9]. The mid-luteal phase has been linked to increased turbulent dynamics and a less stable brain connectome than in the ovulatory phase [5, 9]. The emerging pattern suggests that brain dynamics are lowest during the early follicular phase, increase and peak during ovulation, and fall to intermediate levels during the mid-luteal phase. These fluctuations in brain dynamics are suggested to reflect the influence of hormonal variation across the cycle, particularly changes in estradiol and progesterone concentrations [5, 7, 11–13].

Several analytical approaches have been employed to investigate the impact of the menstrual cycle on brain activity, including independent component analysis [14–19], eigenvector centrality mapping [19, 20], seed-based analysis [17, 19, 20], spectral dynamic causal modeling [21], brain entropy [22], and novel whole-brain dynamic approaches [4, 5, 7, 23]. These studies have significantly advanced our understanding of how female sex hormones influence the functional organization of brain networks, consistently showing menstrual cycle-related changes in the default mode and frontoparietal control networks, and less frequently, changes in the dorsal attention, somatomotor, and limbic networks [4, 5, 7, 14, 16, 18, 19, 21, 22, 24]. However, findings across studies are not always consistent. This variability arises not only from differences in the networks examined but also from the wide range of analytical approaches employed, which can yield divergent results even within the same network or cycle phase. These methodological discrepancies highlight the importance of whole-brain modeling approaches that can robustly capture the complex, distributed effects of hormonal fluctuations on brain dynamics, providing a deeper understanding of the mechanisms underlying menstrual cycle-related brain changes.

In the present study, we apply a thermodynamics-inspired framework to investigate the func-tional hierarchical brain organization throughout the menstrual cycle, based on recent work that quantifies temporal irreversibility in neural signals as a marker of non-equilibrium brain activity [25–27]. Temporal irreversibility reflects the asymmetry in brain signal fluctuations over time and has been proposed as a proxy for the brain’s departure from equilibrium and its underlying func-tional hierarchy. We apply this framework to a large cohort of naturally cycling women scanned in three distinct menstrual phases [7, 21]. We combine it with a whole-brain generative modeling approach, enabling us to move beyond descriptive and correlational measures toward a mechanistic understanding of how endogenous hormonal fluctuations shape brain dynamics. Then, we employ a support vector machine (SVM) classifier on the generative effective connectivity (GEC) matrices, allowing for robust pattern separation in high-dimensional neural data. We hypothesized that fluc-tuations in ovarian hormones across the menstrual cycle modulate the temporal irreversibility of resting-state brain activity, reflecting dynamic shifts in the brain’s functional hierarchy and its dis-tance from equilibrium. We further hypothesized that whole-brain effective connectivity modeling would reveal distinct phase-specific patterns of directional influence between brain areas.

## Results

### Oveview of Methods

To characterize how menstrual cycle-related hormonal fluctuations affect brain dynamics, we used a thermodynamics-inspired approach to quantify temporal irreversibility in resting-state fMRI data. From this, we derived a Non-Reversibility (NR) matrix for each menstrual cycle phase, reflecting the asymmetry in time-shifted interactions across brain regions (see **Figure 1a-e** and Methods). These NR matrices offer a model-free estimate of the system’s functional hierarchy, which we compared across menstrual phases. In addition, we used the NR data to fit a whole-brain generative model, allowing us to construct a generative effective connectivity (GEC) matrix (see **Fig. 1f-h** and Methods). This model-based analysis allows measuring directional influences between brain regions, providing mechanistic insight into how hierarchical brain organization is modulated across the menstrual cycle.

**Figure 1.**
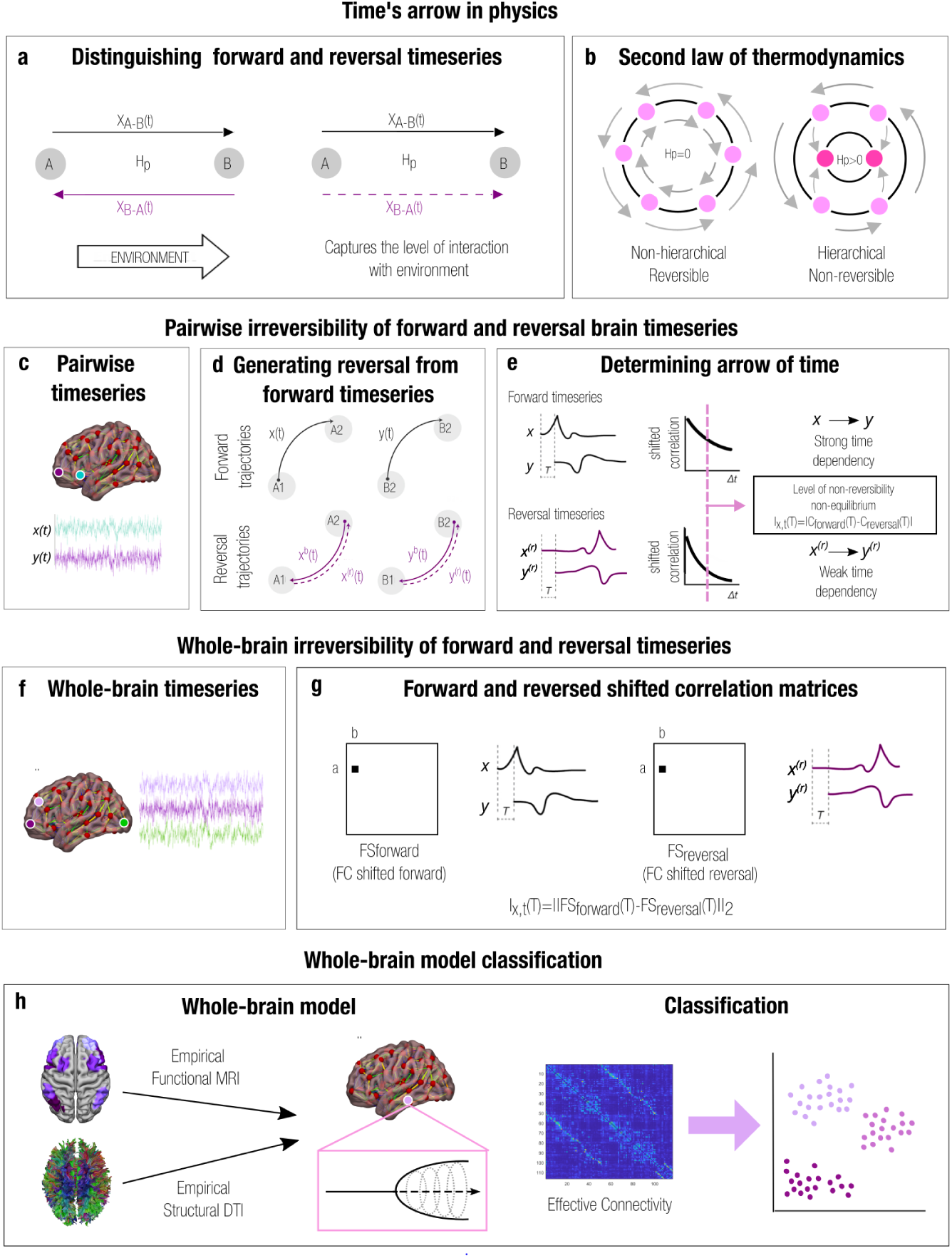
Capturing signatures of brain states by estimating irreversibility. **(a)** Non-equilibrium system with two states A and B and their associated trajectories. Black arrows indicate forward processes, violet arrows indicate backward processes. Whenever the forward and time reversal of backward trajectories differ, the process is considered irreversible. **(b)** Irreversibility can be interpreted as an indirect measure of the hierarchical organization of the system. The panel shows two systems, each with states and their transitions illustrated with circles and arrows, respectively. The left scheme shows a non-hierarchical system, fully reversible over time, with no change in entropy production. The right scheme shows a hierarchical system, with asymmetry in its underlying causal interactions, less reversible over time, with changes in entropy production. **(c)** We can extract the “arrow of time” in brain signals. **(d)** The backward brain signal is constructed by reversing the forward time series in each region. **(e)** For a pair of time series, the shifted correlation illustrates the causal relationships between forward and reversed signals. Reversed time series exhibit a weaker time dependence, as the shifted correlation (as a function of the time shift, Δ*t*) decays more rapidly. **(f)** We applied this framework to the whole brain’s multidimensional time series. **(g)** For the forward and reversed time series, we created two time-shifted correlation matrices (at a given shift time point Δ*t* = *T*). The distance between two matrices determines the level of irreversibility. **(h)** We fitted a whole-brain Hopf model for extracting a generative effective connectivity matrix (*GEC*) linking structural anatomy with functional dynamics based on asymmetric pairwise interactions between brain regions. We then trained a Support Vector Machine classifier to distinguish between menstrual cycle phases using, separately, Functional Connectivity, *GEC*, *GEC* combined with hormonal biomarkers, and hormonal biomarkers alone as input features, separately.

### Demographic data

We analyzed data from 60 healthy young women with regular menstrual cycles, previously described in Hidalgo-Lopez et al. [21]. Participants had a mean age of 25.4 years (range: 18–35) and an average menstrual cycle length of 28 days (range: 23–38). As expected, significant hormonal fluctuations were observed across menstrual phases, consistent with patterns seen in healthy cycling women. Estradiol levels varied significantly, peaking during the pre-ovulatory phase relative to both the early follicular (*β* = 0.329, *SE* = 0.051, *p*_FDR_ *<* 0.001, 95%) and mid-luteal phases (*β* = −0.178, *SE* = 0.051, *p*_FDR_ = 0.003, 95%). Estradiol was also higher in the mid-luteal phase compared to the early follicular phase (*β* = 0.151, *SE* = 0.051, *p*_FDR_ = 0.009, 95%). Progesterone levels showed a clear mid-luteal increase, significantly exceeding levels in the pre-ovulatory (*β* = 116.34, *SE* = 13.05, *p*_FDR_ *<* 0.001) and early follicular phases (*β* = 138.74, *SE* = 13.05, *p*_FDR_ *<* 0.001).

### Hormonal modulation of irreversibility across the whole-brain and resting-state networks

To examine how brain dynamics vary across the menstrual cycle, we applied the model-free thermodynamics framework that quantifies the level of temporal irreversibility in neural activity. In particular, we computed irreversibility by comparing forward and time-reversed time-shifted correlations across brain areas, capturing how past activity in one region predicts future activity in another. This approach was applied to each woman in the early follicular, pre-ovulatory, and mid-luteal phases. Then, we used multilevel Linear Mixed-effects Models (LMMs) to investigate how irreversibility varies across menstrual cycle phases and how it is modulated by individual differences in age and hormone levels, both at the whole-brain level and within resting-state networks. For each network, the irreversibility values of its components were used as dependent variables (see the Statistical analyses section). Note that these values range between 2.16×10^−5^ to 4.80×10^−2^, which constrains the expected order of magnitude of the coefficients. Significant effects are illustrated in **Figure 2**. Our analysis revealed distinct effects of estradiol, progesterone, and their interaction on irreversibility across the whole brain and within specific resting-state networks. Additionally, age was a significant modulator of irreversibility in the default mode and frontoparietal control networks.

**Figure 2:**
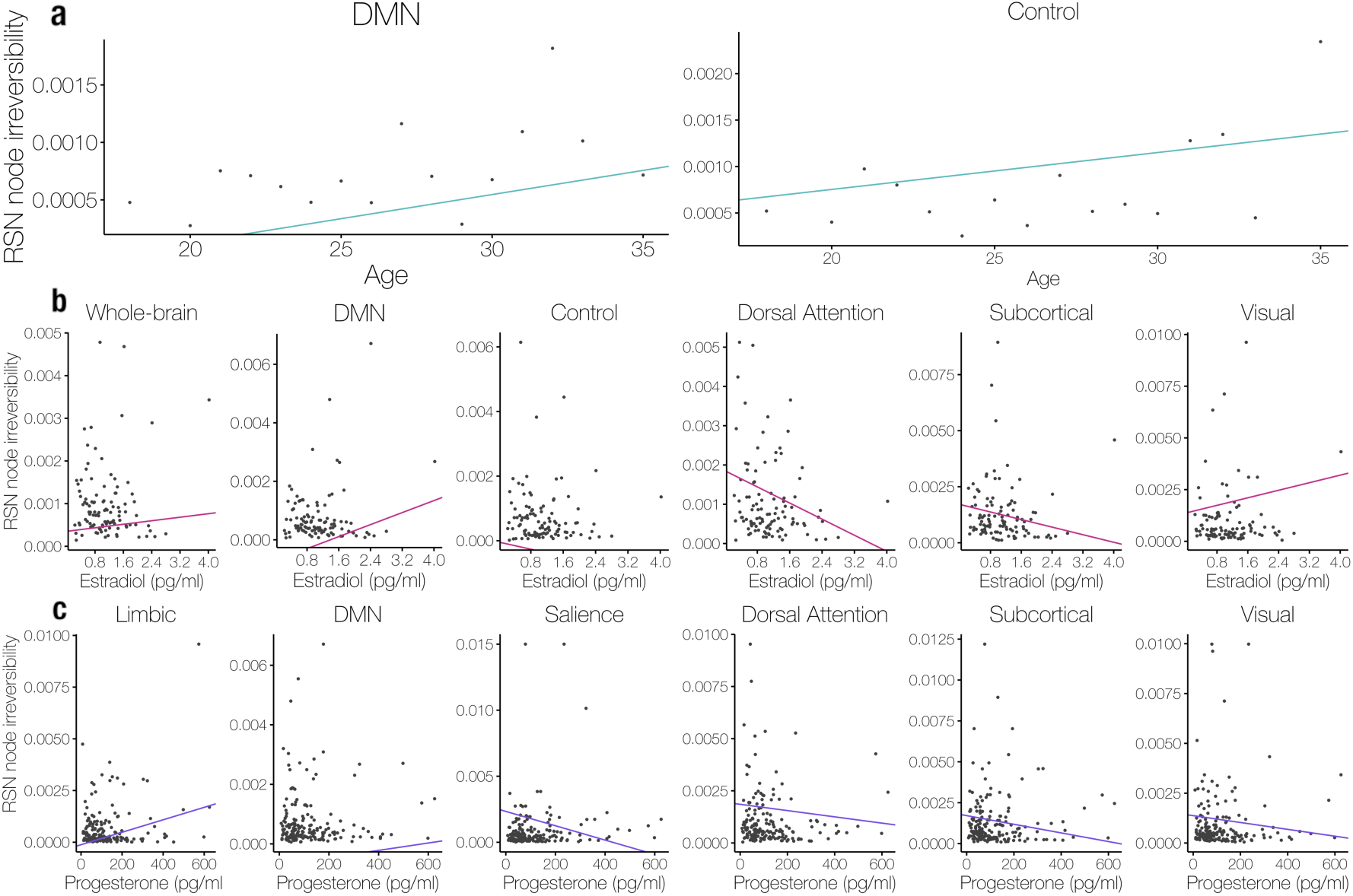
Multilevel linear mixed models of age, estradiol, and progesterone effects on node irreversibil-ity across resting-state brain networks. For each network (subplots), fitted lines represent trajectories predicted by multilevel linear mixed-effects models using data from all participants and corresponding network nodes. Dots indicate the mean node irreversibility at each observed value of the predictor (age or hormone level), averaged across participants and nodes. Lines illustrate the modeled association between predictors and node irreversibility across networks. Higher irreversibility indicates a greater “departure from equilibrium,” i.e., interpreted as increased brain dynamics, energy consumption and information processing. (a) Age (light blue) shows a significant effect on irreversibility in the DMN (*p* = 0.040) and the frontoparietal control network (*p* = 0.038), with increasing age associated with higher irreversibility. (b) Estradiol levels (magenta) significantly affect irreversibility at the whole-brain level (*p* = 0.044), as well as within the DMN (*p <* 0.001), frontoparietal control (*p* = 0.027), dorsal attention (*p* = 0.001), subcortical (*p* = 0.021), and visual networks (*p* = 0.002). Higher estradiol levels are associated with increased irreversibility in the whole-brain, DMN, and visual networks, and decreased irreversibility in the frontoparietal control, dorsal attention, and subcortical networks. (c) Progesterone levels (violet) show significant effects on irreversibility in the limbic (*p <* 0.001), DMN (*p* = 0.002), salience (*p <* 0.001), dorsal attention (*p* = 0.053), subcortical (*p* = 0.002), and visual networks (*p* = 0.013). High progesterone levels are associated with increased irreversibility in the limbic and DMN, and decreased irreversibility in the salience, dorsal attention, subcortical, and visual networks.

At the whole-brain level, both estradiol (*β* = 1.03 × 10^−4^, *SE* = 5.12 × 10^−5^, *p* = 0.044) and the interaction between estradiol and progesterone (*β* = 4.40 × 10^−7^, *SE* = 2.03 × 10^−7^, *p* = 0.029) were statistically significant predictors of irreversibility. These results indicate that elevated estradiol levels are associated with increased temporal asymmetry in brain dynamics, and that this effect is further potentiated when progesterone levels are also high.

For the DMN, age (*β* = 4.20 × 10^−5^, *SE* = 2.05 × 10^−5^, *p* = 0.040), estradiol (*β* = 5.14 × 10^−4^, *SE* = 8.85 × 10^−5^, *p <* 0.001), and progesterone (*β* = 1.24 × 10^−6^, *SE* = 4.13 × 10^−7^, *p* = 0.002) were significant predictors of irreversibility. These results indicate that irreversibility increased with age, estradiol, and progesterone levels. A significant negative interaction was also observed between estradiol and progesterone (*β* = −1.99 × 10^−6^, *SE* = 3.54 × 10^−7^, *p <* 0.001). This interaction suggests that the combined elevation of both hormones leads to a reduced increase in irreversibility, relative to the additive effects of each hormone alone.

For the limbic network, progesterone emerged as a significant predictor of irreversibility (*β* = 3.01 × 10^−6^, *SE* = 8.51 × 10^−7^, *p <* 0.001), indicating that higher progesterone levels are associated with increased irreversibility in the limbic network.

For the dorsal attention network estradiol was a significant negative predictor of irreversibility (estradiol *β* = −5.13×10^−4^, *SE* = 1.61×10^−4^, *p* = 0.001), suggesting that increased estradiol levels were associated with decreased temporal asymmetry. Progesterone showed a marginal negative effect (*β* = −1.50 × 10^−6^, *SE* = 7.76 × 10^−7^, *p* = 0.053), indicating a similar, though less robust, trend. Importantly, a significant positive interaction between estradiol and progesterone was found (*β* = 1.42 × 10^−6^, *SE* = 6.62 × 10^−7^, *p* = 0.032). This interaction suggests that when both hormone levels are elevated (e.g., during the mid-luteal phase), their individual effects on irreversibility are attenuated. In other words, when both estradiol and progesterone are elevated simultaneously, the reduction in irreversibility is less pronounced than when either hormone increases alone.

A similar pattern was observed in the subcortical network, where both estradiol (*β* = −4.22 × 10^−4^, *SE* = 1.83 × 10^−4^, *p* = 0.021) and progesterone (*β* = −2.64 × 10^−6^, *SE* = 8.63 × 10^−7^, *p* = 0.002) were significant negative predictors of irreversibility. These findings suggest that increases in either hormone are associated with reduced directional complexity in subcortical dynamics. Moreover, a significant positive interaction between estradiol and progesterone (*β* = 2.69 × 10^−6^, *SE* = 7.39 × 10^−7^, *p <* 0.001) indicates that their combined presence mitigates the strength of their individual effects. When both hormones increase, the reduction in irreversibility within the subcortical network is less pronounced than when either hormone increases in isolation.

For the salience attention network, progesterone was a significant negative predictor of irreversibility (*β* = −5.43 × 10^−6^, *SE* = 1.00 × 10^−6^, *p <* 0.001), indicating that higher progesterone levels are associated with reduced directional complexity in this network’s dynamics. Additionally, a significant positive interaction between estradiol and progesterone was found (*β* = 5.92 × 10^−6^, *SE* = 8.58 × 10^−7^, *p <* 0.001), suggesting that estradiol attenuates or even reverses the negative effect of progesterone on irreversibility, leading to increased irreversibility.

For the frontoparietal control network, age was a significant positive predictor of irreversibility (*β* = 3.97×10^−5^, *SE* = 1.92×10^−5^, *p* = 0.038), suggesting that older participants exhibited greater directional complexity in this network. In contrast, estradiol was a significant negative predictor (*β* = −2.77 × 10^−4^, *SE* = 1.25 × 10^−4^, *p* = 0.027), indicating that higher estradiol levels were associated with reduced irreversibility in frontoparietal dynamics.

For the visual network, estradiol was a significant positive predictor of irreversibility (*β* = 4.56×10^−4^, *SE* = 1.50×10^−4^, *p* = 0.002), indicating that higher estradiol levels were associated with increased directional complexity in this network. In contrast, progesterone was a significant negative predictor (*β* = −1.72×10^−6^, *SE* = 6.99×10^−7^, *p* = 0.013), suggesting that irreversibility decreased as progesterone levels increase. Additionally, a significant positive interaction between estradiol and progesterone was observed (*β* = 1.45 × 10^−6^, *SE* = 6.00 × 10^−7^, *p* = 0.015), indicating that the estradiol-related increase in irreversibility may neutralize the suppressing effect of progesterone.

Finally, for the somatomotor network, the interaction between estradiol and progesterone was marginally significant (*β* = 8.02 × 10^−7^, *SE* = 4.65 × 10^−6^, *p* = 0.084), suggesting that when estradiol and progesterone are both high, their combined effect may modestly increase irreversibility. Although not below the conventional *p <* 0.05 threshold, this result is worth highlighting as a trend.

### Whole-brain modeling of hierarchical brain dynamics across menstrual cycle phases

For each participant and menstrual phase, we applied the model-based approach to generate a whole-brain model that captures the effective connectivity between brain regions. The model represents each brain region as a Kuramoto oscillator, where local dynamics are influenced by both the intrinsic properties of each region and interactions with other regions. Crucially, the parameters of the model were fit to reproduce the empirical irreversibility measure, which allowed the model to incorporate the directional asymmetry of brain activity. This yielded a generative effective connectivity (*GEC*) matrix, which represents the directional influence between regions. The coupling between regions was optimized iteratively to match the observed brain activity. Specifically, we simulated the BOLD (blood-oxygen-level-dependent) signal, computed Functional Connectivity (*FC*) matrices and time-shifted forward and reverse correlation matrices, and adjusted the model parameters to minimize the difference between empirical and simulated data. This optimization process was repeated until the model accurately reproduced the observed data, resulting in a *GEC* matrix for each participant and menstrual phase. Informed by the irreversibility measure, this whole-brain model was then used to examine how effective connectivity varies across different phases of the menstrual cycle.

### Support vector machine (SVM) classification of menstrual cycle phases using *GEC* matrices

We trained SVM classifiers for the pairwise and tertiary classification of the menstrual cycle phases, using either *GEC*, hormonal biomarkers, *FC*, and *GEC* plus hormonal biomarkers **Figure 3** illustrates the achieved accuracies in each comparison across 1000 folds training the SVM with *GEC* matrices, hormone levels alone, *FC* matrices, and *GEC* + hormone levels.

**Figure 3:**
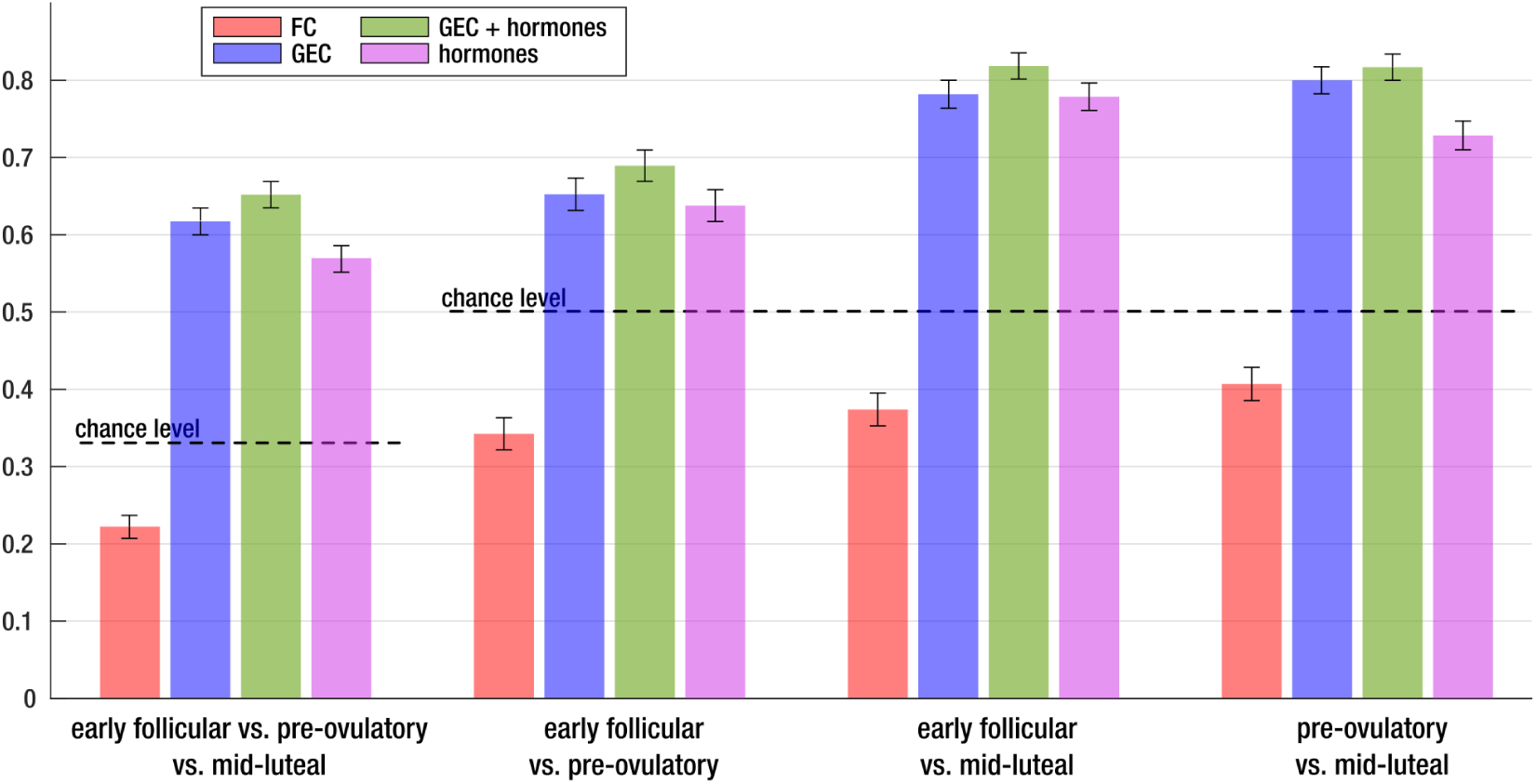
Classification accuracy of menstrual cycle phase using dynamic, static connectivity features and hormonal levels. Support Vector Machine (SVM) classification performance across 1000 folds using a leave-one-out cross-validation scheme. Models are trained using four sets of input features: functional connectivity (*FC*) matrices, generative effective connectivity (*GEC*) matrices, *GEC* combined with hormonal biomarkers (*GEC + hormones*), and hormone levels alone. Bars indicate mean classification accuracy; whiskers represent 95% confidence intervals. Dashed lines indicate chance-level performance. The *FC* model yields the lowest performance across all comparisons (three-phase: 22%; early follicular vs. pre-ovulatory: 34%; early follicular vs. mid-luteal: 37%; pre-ovulatory vs. mid-luteal: 41%). The *GEC* model performs substantially better (three-phase: 62%; early follicular vs. pre-ovulatory: 65%; early follicular vs. mid-luteal: 78%; pre-ovulatory vs. mid-luteal: 80%). The *GEC + hormones* model provides the best overall performance (three-phase: 65%; early follicular vs. pre-ovulatory: 69%; early follicular vs. mid-luteal: 82%; pre-ovulatory vs. mid-luteal: 82%). The hormones-only model shows intermediate accuracy (three-phase: 57%; early follicular vs. pre-ovulatory: 64%; early follicular vs. mid-luteal: 78%; pre-ovulatory vs. mid-luteal: 73%). These results highlight the unique contribution of dynamic, hormone-sensitive connectivity patterns captured by the *GEC* + *hormones* model in discriminating between menstrual cycle phases.

The classification of the *GEC* matrices yielded an accuracy of 62% for the tree-phase (tertiary) classification, with pairwise accuracies of 65% for the early follicular vs. preovulatory, 78% for the early follicular vs. midluteal, and 80% for the preovulatory vs. midluteal phases. Importantly, these models consistently outperformed hormone-only classifiers (57% tertiary; 64% early follicular vs. pre-ovulatory, 78% early follicular vs. mid-luteal, and 73% pre-ovulatory vs. mid-luteal), indicating that *GEC* matrices provide richer information than hormonal measures alone. By contrast, functional connectivity (FC) performed substantially worse (22% tertiary; 34–41% pairwise). Together, these results establish GEC as a more informative biomarker of menstrual phases than hormone levels or FC alone.

As a complementary analysis, GEC matrices were concatenated with hormone features, modestly increasing classification accuracy (from 65% tertiary to up to 82% in pairwise comparisons). While combining GEC with hormones yielded higher accuracies, interpretation should be cautious since hormone measurements were also used to define phase labels. Nonetheless, the improvement suggests potential value in multimodal integration for future research employing independent phase definitions.

In addition, we visualize which brain areas exhibited the most pronounced changes across menstrual cycle phases. For this purpose, we summarized each subject’s *GEC* matrix by computing the average of each row and column, yielding a single value per node. We then calculated the mean difference in these node-wise values between phases, normalized by the sum of their variances, and projected the results onto a 3D brain rendering (see **Figure 4**). The resulting maps reveal that changes in effective connectivity are distributed across multiple functional networks and are spatially widespread rather than confined to specific regions. These differences appear most pronounced from early follicular to mid-luteal and from pre-ovulatory to mid-luteal than from early follicular to pre-ovulatory.

**Figure 4:**
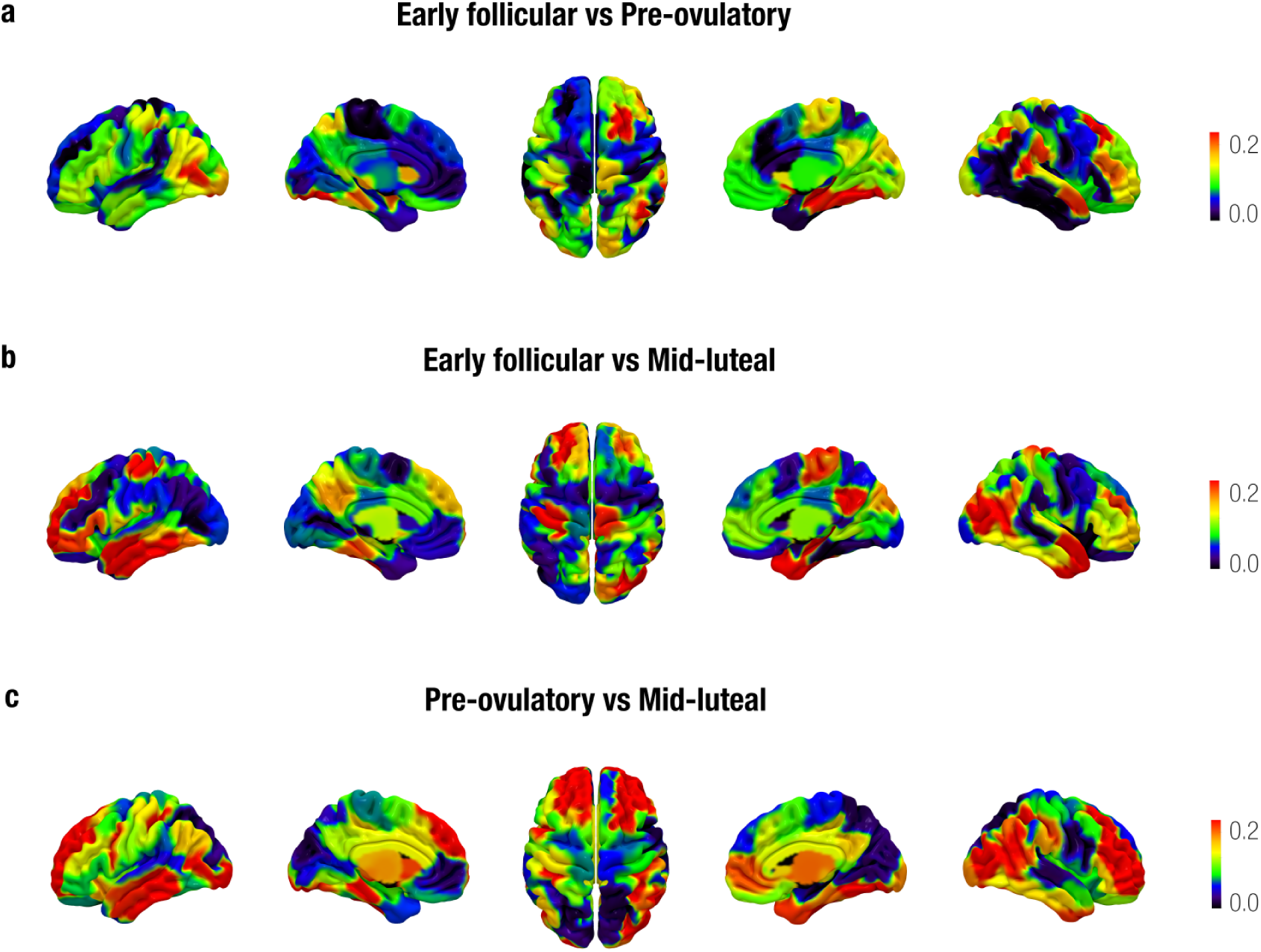
**Brain renderings illustrating differences in generative effective connectivity (***GEC***) across menstrual cycle phases.** Panels show contrasts between: (a) early follicular vs. pre-ovulatory, (b) early follicular vs. mid-luteal, and (c) pre-ovulatory vs. mid-luteal phases. For each participant, the *GEC* matrix was transformed into a one-dimensional array by summing the average of each row and column. Group-level differences were computed by subtracting the phase-wise means and normalizing them by the sum of their variances. Color bars indicate the magnitude of these differences, with blue representing little to no difference and red indicating the greatest differences. The brain renderings show that changes across menstrual cycle phases involve several functional networks rather than being localized. The magnitude of these differences is greater when comparing the early follicular phase to the mid-luteal phase and the pre-ovulatory phase to the mid-luteal phase than when comparing the early follicular phase to the pre-ovulatory phase.

Furthermore, we examined the ten most important node interactions (entries in the *GEC* ma-trix) for the training of the SVM classifiers. These top-ranked interactions are visualised in **Figure 5**, where red arrows represent homotopic interactions (connections between anatomically corre-sponding regions in opposite hemispheres) and black arrows indicate non-homotopic interactions (connections between regions that are not anatomically mirrored across hemispheres). The complete list of interactions for the tertiary and each pairwise comparison are provided in **Tables 1,2,3** and **4**, where interactions are ranked in descending order of importance. For each node pair, we also report regions names, spatial coordinates and associated resting-state network labels to aid interpretation. Across all comparisons, the DMN, dorsal attention, and visual networks consistently featured among the top-ranked interactions. The early-follicular vs. pre-ovulatory contrast closely mirrored the tertiary three-phase classification, with a homotopic dorsal-attention interaction as the single most influential feature, followed by multiple DMN interactions. In the early-follicular vs. mid-luteal comparison, homotopic connections in the visual network predominated. Finally, distinguishing pre-ovulatory from mid-luteal phases again highlighted the visual network and DMN, alongside two homotopic interactions within subcortical regions.

**Figure 5:**
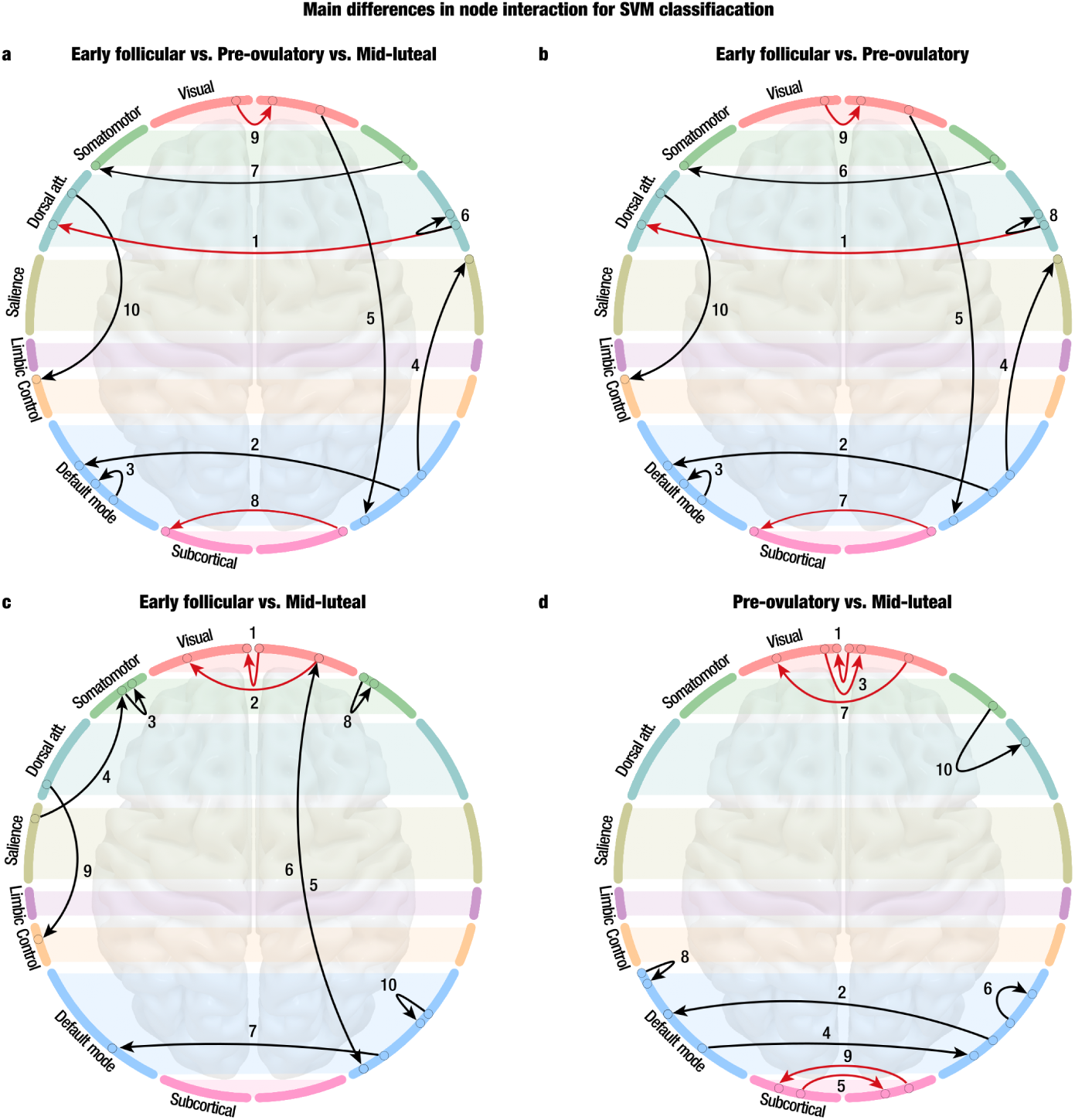
Top 10 most influential node interactions for the Support Vector Machine classification of menstrual cycle phases. Graph representations illustrate the 116 brain regions (nodes), arranged along the perimeter of each circle, with the left and right hemispheres positioned on the corresponding sides. Nodes are color-coded by their respective resting-state networks, indicated by horizontal divisions. Circles highlight the nodes involved in the top 10 most influential interactions contributing to classification accuracy, with filled colors denoting network identity. Directed arrows represent connections between node pairs: red arrows denote homotopic interactions (i.e., connections between anatomically corresponding regions in opposite hemispheres), while black arrows represent non-homotopic interactions (i.e., connections between non-mirrored regions). Numerical ranks (1–10) reflect the relative importance of each interaction in the classification model. **(a) Early follicular vs. Pre-ovulatory vs. Mid-luteal.** The most important feature is a homotopic interaction within the dorsal attention network, followed by several interactions involving the DMN. **(b) Early follicular vs. Pre-ovulatory.** The top-ranked features overlap with those from the three-phase classification, though their relative order differs slightly. **(c) Early follicular vs. Mid-luteal.** Homotopic interactions within the visual network are most influential, followed by interactions involving the somatomotor network. **(d) Pre-ovulatory vs. Mid-luteal.** Homotopic interactions within the visual network again play a prominent role, along with several connections involving the DMN. Notably, two homotopic interactions within subcortical regions also emerge among the most important features.

**Table 1:** Top 10 most influential node interactions for SVM classification between the early follicular, pre-ovulatory, and mid-luteal phases. Each row lists a pair of interacting nodes, along with their spatial coordinates and corresponding resting-state networks. Interactions are ranked in descending order of importance based on their contribution to the classification model. RSN: resting-state network, Dorsal: dorsal attention network, DMN: default mode network, Salience: salience network, SomMot: somatomotor network, Subco: subcortical network, Visual: visual network, Control: prefrontal control network.

| Top Node Interactions: Early Follicular vs. Pre-ovulatory vs. Mid-luteal |  |  |  |  |  |  |  |  |  |
| --- | --- | --- | --- | --- | --- | --- | --- | --- | --- |
| Node In |  |  |  |  | Node Out |  |  |  |  |
| RSN | Node | x | y | z | RSN | Node | x | y | z |
| Dorsal | *LH_DorsAttn_Post_6* | 0 | 118 | 19 | Dorsal | *RH_DorsAttn_Post_5* | 4 | 119 | 19 |
| DMN | *LH_Default_PFC_1* | 205 | 63 | 76 | DMN | *RH_Default_PFCv_2* | 209 | 61 | 82 |
| DMN | *LH_Default_PFC_3* | 205 | 63 | 79 | DMN | *LH_Default_PFC_5* | 205 | 63 | 81 |
| Saliency | *RH_SalVentAttn_TempOccPar_1* | 201 | 58 | 252 | DMN | *RH_Default_Temp_3* | 209 | 61 | 81 |
| DMN | *RH_Default_pCunPCC_1* | 208 | 62 | 78 | Visual | *RH_Vis_6* | 124 | 18 | 139 |
| Dorsal | *RH_DorsAttn_Post_4* | 4 | 119 | 18 | Dorsal | *RH_DorsAttn_Post_5* | 4 | 119 | 19 |
| SomMot | *LH_SomMot_6* | 70 | 130 | 185 | SomMot | *RH_SomMot_8* | 74 | 130 | 187 |
| Subco | *HIP_lh* | 171 | 243 | 85 | Subco | *HIP_rh* | 171 | 243 | 85 |
| Visual | *RH_Vis_2* | 124 | 18 | 134 | Visual | *LH_Vis_2* | 120 | 18 | 132 |
| Control | *LH_Cont_Par_1* | 230 | 148 | 35 | Dorsal | *LH_DorsAttn_Post_3* | 0 | 118 | 16 |

**Table 2:**
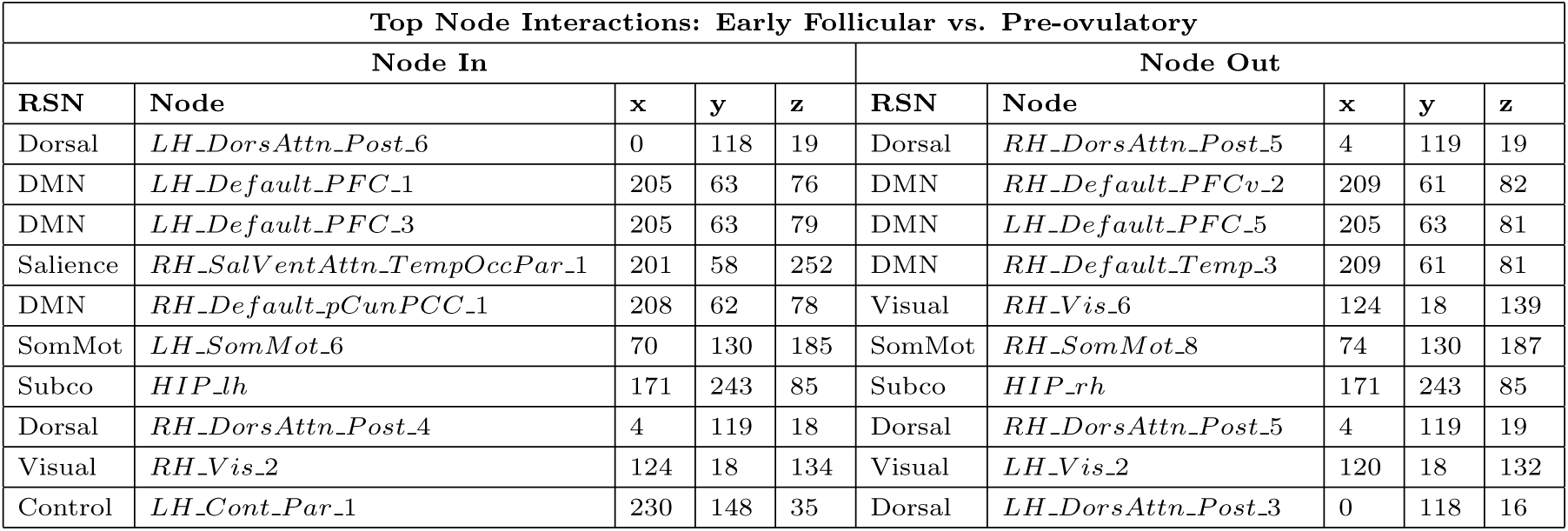
Top 10 most influential node interactions for the SVM classification between the early follicular and pre-ovulatory phases. Each row lists a pair of interacting nodes, along with their spatial coordinates and corresponding resting-state networks. Interactions are ranked in descending order of importance based on their contribution to the classification model.

**Table 3:**
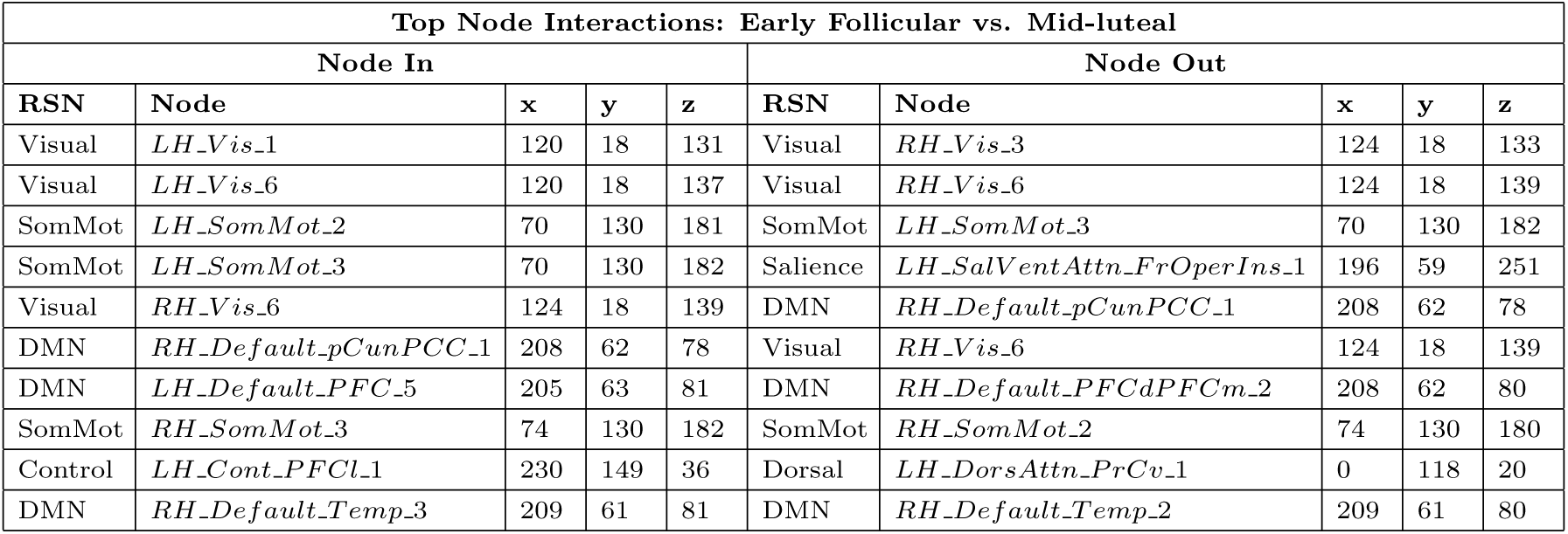
Top 10 most influential node interactions for the SVM classification between the early follicular and mid-luteal phases. Each row lists a pair of interacting nodes, along with their spatial coordinates and corresponding resting-state networks. Interactions are ranked in descending order of importance based on their contribution to the classification model.

**Table 4:** Top 10 most influential node interactions for the SVM classification between the pre-ovulatory and mid-luteal phases. Each row lists a pair of interacting nodes, along with their spatial coordinates and corresponding resting-state networks. Interactions are ranked in descending order of importance based on their contribution to the classification model.

| Top Node Interactions: Pre-ovulatory vs. Mid-luteal |  |  |  |  |  |  |  |  |  |
| --- | --- | --- | --- | --- | --- | --- | --- | --- | --- |
| Node In |  |  |  |  | Node Out |  |  |  |  |
| RSN | Node | x | y | z | RSN | Node | x | y | z |
| Visual | *LH_Vis_1* | 120 | 18 | 131 | Visual | *RH_Vis_1* | 124 | 18 | 133 |
| DMN | *LH_Default_PFC_1* | 205 | 63 | 76 | DMN | *RH_Default_PFCv_2* | 209 | 61 | 82 |
| Visual | *RH_Vis_2* | 124 | 18 | 134 | Visual | *LH_Vis_2* | 120 | 18 | 132 |
| DMN | *RH_Default_PFCdPFCm_2* | 208 | 62 | 80 | DMN | *LH_Default_PFC_5* | 205 | 63 | 81 |
| Subco | *NAc - rh* | 184 | 210 | 157 | Subco | *NAc - lh* | 184 | 210 | 157 |
| DMN | *RH_Default_Par_1* | 209 | 62 | 78 | DMN | *RH_Default_Temp_3* | 209 | 61 | 81 |
| Visual | *LH_Vis_6* | 120 | 18 | 137 | Visual | *RH_Vis_6* | 124 | 18 | 139 |
| DMN | *LH_Default_Temp_2* | 205 | 62 | 81 | DMN | *LH_Default_Temp_1* | 205 | 62 | 80 |
| Subco | *pTHA - lh* | 127 | 130 | 120 | Subco | *pTHA - rh* | 127 | 130 | 120 |
| Dorsal | *RH_DorsAttn_Post_2* | 4 | 119 | 16 | SomMot | *RH_SomMot_6* | 74 | 130 | 185 |

## Discussion

This study employed a thermodynamics-inspired framework to characterize how the brain’s dynamic and hierarchical organization undergoes significant modulation across the menstrual cycle, with a focus on temporal irreversibility and generative effective connectivity (GEC). First, age and ovarian hormones (estradiol and progesterone) significantly and distinctly modulate the brain’s functional hierarchical organization, reflected in changes of temporal irreversibility among large-scale networks. Second, generative effective connectivity (GEC) matrices were identified as robust biomarkers for menstrual phase classification. Model-based analyses demonstrate that GEC, particularly when combined with hormonal measures, achieves superior classification accuracies (up to 82%), significantly outperforming models based solely on traditional functional connectivity (FC) or hormonal biomarkers. Finally, feature importance analysis highlighted homotopic inter-hemispheric connections, predominantly within the DMN, dorsal attention, salience, and visual networks, as the most discriminative features distinguishing menstrual phases. These results provide novel and mechanistic insights into how directed causal brain interactions reorganize throughout menstrual cycle phases, serving as sensitive indicators of hormone-brain interactions.

Model-free analyses demonstrated a network-specific association between ovarian hormones and the temporal irreversibility of brain dynamics. Temporal irreversibility reflects the inherent asymmetry of brain signal fluctuations over time, serving as a marker of the brain’s departure from equilibrium and its underlying hierarchical and directional organization. First, elevated estradiol levels, characteristic of the pre-ovulatory phase, were consistently linked to a more hierarchical and temporally asymmetric organization of large-scale brain networks, reflecting an increased”arrow of time” in neural activity. This finding aligns with prior evidence highlighting estradiol’s pivotal role in widespread functional and structural brain reorganization [4, 7, 24]. Crucially, estradiol’s impact on irreversibility exhibited marked network specificity: positive correlations were observed primarily at the whole-brain level and within DMN and visual networks, while negative correlations emerged in the prefrontal control, dorsal attention, and subcortical networks. These differential effects suggest that estradiol optimizes specific neural circuits for varying cognitive and sensory demands across the cycle. For instance, increased irreversibility in DMN might signify enhanced self-referential processing, while decreased irreversibility in dorsal attention networks could indicate a more efficient or streamlined attentional state. Second, higher progesterone concentrations were associated with increased irreversibility in limbic and DMN networks, yet decreased irreversibility in salience, dorsal attention, subcortical, and visual networks. While some aspects align with studies linking progesterone to enhanced metastability within comparable networks [7] and increased eigenvector centrality in the dorsolateral prefrontal cortex alongside strengthened hippocampal connectivity [20], its overall modulatory effects remain heterogeneous across studies. This is consistent with some reports of reduced intrinsic DMN connectivity, increased low-frequency signal amplitude in the caudate [19], heightened within-DMN effective connectivity, and diminished connectivity in salience and executive control networks [21]. Furthermore, structural neuroimaging investigations also reveal progesterone-associated volumetric brain changes [28], although contradictory findings have been reported [29]. Contrasting these local network-level effects, evidence of reduced whole-brain coherence [24] and decreased entropy in frontoparietal and limbic regions [22] suggests a reduction in global complexity. This combined evidence suggests that progesterone may selectively enhance localized, directional neural signaling within specific networks while concurrently reducing more global or integrative aspects of brain dynamics. Additionally, a notable, though marginally significant, interaction between estradiol and progesterone in the somatomotor network indicates a synergistic enhancement of directional complexity. This aligns with previous reports associating progesterone with increased eigenvector centrality in the sensorimotor cortex [20], estradiol-related increases in sensorimotor coherence [24], and progesterone-associated reductions in metastability throughout the menstrual cycle [7]. Collectively, these results indicate an increased sensitivity of the somatomotor system to coordinated ovarian hormone modulation, potentially facilitating adaptive sensorimotor regulation. Finally, the observed age-related increases in temporal irreversibility within the DMN and frontoparietal control networks are consistent with neuroimaging studies showing peak dynamical complexity in middle age, followed by a decline after menopause [7, 30]. This reinforces the interconnected effects of reproductive aging and hormones on large-scale neural dynamics.

In the model-based analyses, generative effective connectivity (*GEC*) matrices were employed to capture directed, causal interactions across the whole-brain network for each menstrual cycle phase. Derived from a mechanistic generative model of brain dynamics [27], these *GEC* matrices provide asymmetric, directional connectivity information. Unlike FC, which quantifies undirected statistical dependencies, *GEC* captures the influence that one brain region exerts over another, thereby offering a more accurate representation of the communication between brain regions. Generative effective connectivity (GEC) matrices proved to be the most informative feature set for menstrual phase classification. Compared to functional connectivity (FC), which performed near chance levels, GEC provided substantially higher accuracy in both tertiary and pairwise classifications, underscoring the superior sensitivity of directed, causal interactions in capturing hormone-driven reorganization of brain networks. Hormone-only models performed moderately well but consistently under-performed relative to GEC, suggesting that connectivity-based features convey richer information than endocrine markers alone. Notably, when GEC features were combined with hormonal profiles, classification improved further, surpassing 80% accuracy in some pairwise comparisons. While such multimodal integration highlights the potential of combining complementary information, caution is warranted given that hormone levels were also used to define menstrual cycle phases. Overall, these findings position GEC as a promising avenue for developing robust, mechanistically grounded biomarkers of hormone-related brain state transitions. These findings align with those of Hidalgo-Lopez and colleagues [21], who applied spectral dynamic causal modeling to investigate changes in effective connectivity within the default mode, salience, and executive control networks. Using a leave-one-out cross-validation approach, they identified key effective connections whose modulation enabled above-chance prediction of menstrual phases, indicating hormone-related network reconfiguration. Together, these complementary approaches highlight effective connectivity as a sensitive feature space for decoding hormone-brain interactions, suggesting that focusing on causal connections within key large-scale networks can enhance individualized prediction of menstrual phases. Beyond menstrual cycle classification, machine learning techniques have been employed to distinguish individuals with and without primary dysmenorrhea [31, 32], characterize reproductive stages across the adult lifespan [30], and predict sex and gender in children and preadolescents using resting-state functional MRI data [33, 34]. Collectively, these studies underscore the utility of machine learning methods for detecting hormone-sensitive and sex-related brain signatures, with significant implications for advancing women’s brain health research and clinical applications.

Finally, we identified the most significant features that distinguished the menstrual cycle phases. The feature importance analysis for the tertiary menstrual cycle phase classification identified directed effective connectivity patterns as robust biomarkers for menstrual cycle phase classification, with homotopic interhemispheric connections within the DMN, dorsal attention, and visual networks being especially discriminative. These bilateral coordinated dynamics corroborate prior neuroimaging evidence showing menstrual phase–dependent modulation of these networks, traditionally observed via undirected functional connectivity measures [5, 7, 21, 24]. The mechanistic modeling approach, leveraging GEC matrices, advances this understanding by identifying directional, phasespecific effective connectivity as crucial markers of cycle transitions. Pairwise analyses revealed phase-specific connectivity changes consistent with known hormonal fluctuations. The early follicular to pre-ovulatory transition prominently featured homotopic interhemispheric connections within the dorsal attention network, alongside cross-network directed interactions between the DMN and salience network involving hippocampal nodes. This pattern aligns with studies reporting that estradiol peaks are associated with enhanced modulation of attentional and memory systems [7, 20, 35], reflecting both bilateral coordination within attentional circuits and asymmetric causal influences across large-scale neural networks. The early follicular to mid-luteal contrast was characterized by pronounced homotopic visual network connectivity coupled with significant non-homotopic somato-motor interactions, as well as notable modulations in salience and frontoparietal control networks. These findings are consistent with progesterone’s established effects on sensorimotor and visual system dynamics [7, 20, 23, 36], highlighting complex hierarchical reorganization of sensory and cognitive control circuits across menstrual phases. Finally, the comparison between pre-ovulatory and mid-luteal phases expanded these findings by revealing widespread bilateral homotopic effective connectivity within the DMN and visual cortices, alongside substantial involvement of subcortical regions, including the nucleus accumbens and thalamus. This pattern supports prior reports showing the sensitivity of the basal ganglia and hippocampus to ovarian hormones [37, 38], emphasizing the integrative role of cortico-subcortical loops in hormonally driven brain reconfiguration. Overall, these results demonstrate that effective menstrual cycle phase decoding relies on distributed, hierarchically organized brain networks and richly interacting cortical and subcortical circuits, consistent with hormone-sensitive modulation of cognitive, attentional, affective, and sensorimotor processes [5, 7, 20, 23, 24, 36, 38, 39]. Future research should explore the clinical and behavioral implications of these directional connectivity biomarkers, which may hold promise for precision diagnostics for menstrual cycle–related brain disorders.

Several methodological limitations warrant consideration. First, our analyses focused on estradiol and progesterone concentrations; however, other key hormones involved in menstrual cycle regulation, such as luteinizing hormone and follicle-stimulating hormone, may provide additional insight. Second, although menstrual phase definitions adhered to established criteria, inter-individual variability in cycle length and hormone timing may have introduced noise into phase assignment, potentially impacting classification accuracy and network characterization. Addressing these challenges will require longitudinal, within-subject designs with dense sampling protocols that combine daily neuroimaging with comprehensive, multi-hormonal profiling throughout the menstrual cycle [24, 40]. Third, we employed a well-defined resting-state parcellation to obtain the time series; however, network topology and connectivity patterns can vary depending on the chosen brain atlas, which can influence the results [41]. Fourth, the functional MRI data were collected with a repetition time (TR) of 2.25 seconds. This temporal resolution limitation may reduce sensitivity to rapid neural fluctuations and affect the accuracy of effective connectivity estimates. Finally, consistent with recent calls in neuroendocrinology and female mental health research [42, 43], future studies should include larger, demographically diverse, and adequately powered cohorts, employing longitudinal and multimodal designs. Such approaches are essential for capturing the complex individual variability in hormone-brain interactions and ensuring the generalizability and robustness of findings across populations.

In summary, our study demonstrates that generative effective connectivity (GEC) matrices provide a sensitive and physiologically grounded feature set for distinguishing menstrual cycle phases. Using machine learning classifiers, GEC-based models achieved substantially higher accuracy in classifying both tertiary and pairwise menstrual cycle phases compared to conventional functional connectivity and hormone-only approaches. Key brain networks—including the DMN, dorsal attention, visual, and subcortical—showed phase-specific directed interactions that underpin hormonally driven brain reorganization. These findings underscore the mechanistic relevance of directed causal interactions in brain networks as biomarkers of hormone-related brain state transitions, establishing a foundation for more precise approaches to decode female brain function across the menstrual cycle. Importantly, these findings may have clinical relevance for characterizing menstrual cycle–related disorders such as premenstrual dysphoric disorder and primary dysmenorrhea. This approach could enhance diagnostic precision, facilitate individualized monitoring, and inform the development of targeted therapeutic interventions that address brain–hormone interactions.

## Methods

### Participants

A total of 60 healthy young women were selected from a dataset described in Hidalgo-Lopez et al. [21]. Their age ranged between 18 and 35 years, and their menstrual cycles lasted between 21 and 35 days with an intercycle variability of less than 7 days. Participants met the following inclusion criteria: (1) no use of hormonal contraceptives in the 6 months prior to the study, (2) no history of neurological, psychiatric, or endocrine disorders, and (3) no current medication use. Each participant was scheduled for three appointments, one during each of the following phases: pre-ovulatory, early follicular, and mid-luteal. These corresponded to 2-3 days before the expected onset of ovulation, 1-7 days after the onset of current menses, and 3 days after ovulation to 3 days before the expected onset of the next menstruation. Appointments were counterbalanced to minimize potential bias. Ovulatory urine tests (Pregnafix ®) confirmed the pre-ovulatory phase, and participants later confirmed the onset of their next menstruation. The cycle duration was calculated based on participants’ self-reported onset dates of the last three cycles. All participants provided written informed consent. The study was approved by the University of Salzburg Ethics Committee in accordance with the Declaration of Helsinki.

### Hormone analysis

Salimetrics salivary estradiol and progesterone ELISAs were used to measure hormone levels. Samples of saliva were first collected with the passive drool method, stored at-20°C and centrifuged at 3000 rpm twice (15 min and 10 min, respectively) to remove solid particles. Duplicates of samples were obtained and the ones presenting more than 25% of variation were re-analyzed. LMMs assessed differences in estradiol and progesterone levels across menstrual cycle phases (see Statistical analyses section).

### MRI data acquisition

A Siemens Magnetom TIM Trio 3T scanner was used to obtain the MRI scans. Resting-state fMRI was performed using a T2^∗^ weighted gradient echo-planar (EPI) sequence (volumes= 244; number of slices = 36; TE = 30 ms; TR = 2250 ms; flip angle (FA) 70°; slice thickness = 3.0mm; matrix 192 × 192; FOV 192 mm; in-plane resolution 2.6 × 2.6 mm). Structural T1-weighted images were acquired with 160 sagittal slices (TE=291 ms; TR=2300 ms; TI delay=900 ms; (FA) 9°; slice thickness = 1 mm; FOV 256 × 256 mm). Participants were asked to relax, close their eyes, and allow their thoughts to flow freely. Diffusion-weighted imaging (DWI) data were obtained with a dual spin-echo diffusion tensor imaging (DTI) sequence, consisting of 60 contiguous axial slices, with isotropic voxel size 2 mm, no gap, matrix sizes 118 x 118, TE 92 ms and FOV 236 mm. Diffusion was recorded with 64 optimal non-collinear diffusion directions with a b value of 1,500 s/mm2 interleaved with 9 non-diffusion b0 images. A frequency-selective fat saturation pulse was used to minimize artifacts.

### Resting-state fMRI preprocessing

The first six volumes of each participant’s scan were discarded, and functional images were despiked using the AFNI 3D-despiking algorithm [44–46]. Despiked images were then preprocessed following standard procedures in SPM12 (www.fil.ion.ucl.ac.uk/spm). The segmentation of the structural images was performed using CAT12. Then, the resulting images were cleaned using the ICA-AROMA algorithm in FSL to remove artifactual components in a non-aggressive manner [47, 48]. Finally, the blood oxygenation level-dependent (BOLD) time series were filtered with a second-order Butterworth filter in the band 0.04–0.08 Hz and extracted according to a well-defined resting-state cortical atlas of 100 nodes, belonging to 7 resting-state networks [49], and a subcortical atlas of 16 nodes [50].

### Probabilistic tractography analysis

A whole-brain structural connectivity matrix (SC) template was computed averaging SC matrices from 40 unrelated female participants, from a cohort previously described in [51]. All SC matrices were computed in native MRI diffusion space using the same parcellation described above. Analyses were conducted using the FMRIB’s Diffusion Toolbox (FDT) in FMRIB Software Library (www.fmrib.ox.ac.uk/fsl), as successfully applied in similar studies [52–54]. In brief, DICOM images were converted to NIfTI format using dcm2nii (www.nitrc.org/projects/dcm2nii). B0 images were co-registered to T1-weighted images in native space using FLIRT [55], and the co-registered T1 images were further aligned to standard space using FLIRT and FNIRT [55, 56]. The transformations were inverted and applied to warp the MNI-space atlas into each subject’s native diffusion space using nearest-neighbor interpolation. Diffusion-weighted images were then analyzed using the FDT. Brain extraction was performed using the Brain Extraction Tool (BET) [57], and eddy current distortions and head motion were corrected using the eddy correct tool [58]. Gradient matrices were re-oriented to account for subject motion [59]. Crossing fibers were modeled using BEDPOSTX, which estimates the probability of multiple fiber orientations, enhancing sensitivity to non-dominant fiber populations [60, 61]. Finally, probabilistic tractography was performed in native diffusion space using PROBTRACKX following standard procedures.

### Assessing irreversibility in timeseries

The level of irreversibility was computed by assessing the degree of asymmetry in the temporal evolution of brain activity [26]. Specifically, given two time series *x*(*t*) and *y*(*t*), we assumed that *x*(*t*) evolved from an initial state *A*_1_ to a final state *A*_2_, and that *y*(*t*) evolved from an initial state *B*_1_ to a final state *B*_2_. Their reversed versions were obtained by flipping the time series (**Figure 1 d**). More specifically, *x*^(*r*)^(*t*), the reversed version of *x*(*t*), was generated by inverting the temporal order, evolving from state *A*_2_ to *A*_1_. The same procedure was applied to *y*(*t*) to obtain *y*^(*r*)^(*t*). Time-shifted correlation was then used to estimate the causal dependency between two time series (**Figure 1e**). For the forward evolution, we computed the time-shifted correlation as:

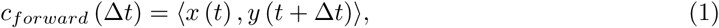

or the reversed evolution as:

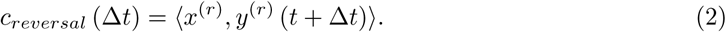

We quantified the pairwise level of irreversibility (i.e., temporal asymmetry), which captures the “arrow of time,” as the absolute difference between forward and reversal correlations at a time lag of Δ*t* = *T*:

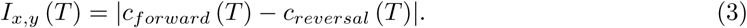

We selected *T* = 2 (i.e., a shift of two time points) after optimization to maximize statistical significance. For multidimensional data, we extended this approach by computing forward and reversal time-shifted correlation matrices (see **Figure 1 g**). Let *x_i_*(*t*) denote the *i*-th time series and *x*^(*r*)^(*t*) its reversed counterpart. Functional causal dependencies (***F S***) were then calculated as:

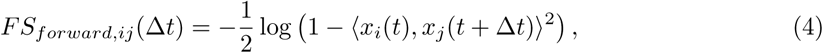

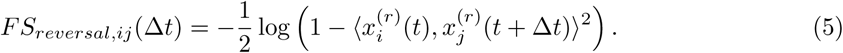

The forward and reversal time-shifted correlation matrices *FS* were expressed in terms of mutual information to ensure positive values. We obtained pairwise irreversibility by computing the squared difference between forward and reversal matrices at Δ*t* = *T*:

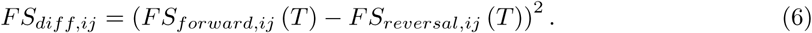

Finally, we calculated the total irreversibility for each brain area (or node) as the sum of the mean across across rows plus the mean across columns of ***F S***_diff_.

### Whole-brain modeling

We fit individualized whole-brain models, which yielded a generative effective connectivity ma-trix for each subject in each phase as described in [27]. The model represents each brain region as a Kuramoto oscillator, with local dynamics governed by the normal form of a supercritical Hopf bifurcation. In complex coordinates, each node *j* is described by:

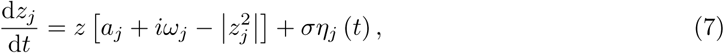

where

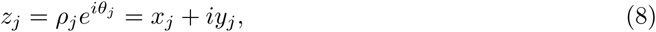

and *η_j_*(*t*) denotes Gaussian noise with standard deviation *σ* = 0.02. This system undergoes a supercritical bifurcation at *a_j_* = 0, with a stable fixed point at *z_j_* = 0 for *a_j_ <* 0, and a stable limit cycle with oscillation frequency *f_j_* = *<u>^ωj^</u>*for *a_j_ >* 0. Inserting Equation 8 into Equation 7 and separating real and imaginary components in equations 9 and 10 respectively, gives:

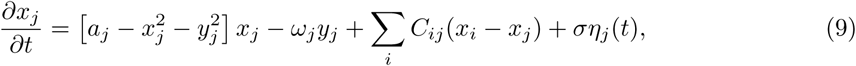

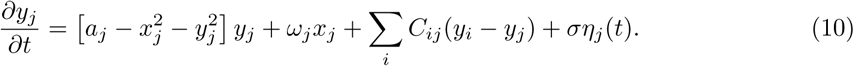

We set the bifurcation parameter to *a_j_* = −0.02 following prior work [62]. Intrinsic frequencies *ω_j_* were computed by averaging the peak frequency of the bandpass-filtered BOLD signal from each of the 116 brain regions.

The *GEC* matrix was obtained by optimizing the coupling matrix ***C*** to reproduce each subject’s dynamics, including the empirical functional connectivity (***F C***) and forward/reversal shifted-correlation matrices (***F S***). Specifically, the *GEC* matrix was derived by optimizing the coupling matrix ***C***, initially informed by the structural connectivity matrix (***SC***). Each model was iteratively updated over 1500 optimization steps and the distance between the empirical and modeled ***F C*** and the forward and reversed ***F S*** was calculated iteratively to update the *GEC* matrix. In each iteration, BOLD data were simulated 100 times, given the simulated BOLD data, the functional connectivity ***F C****^model^*, forward ***F S****^model^* (Δ*t*) and reversal ***F S****^model^* (Δ*t*) shifted correlation matrices were computed (equations 4 and 5). These matrices were averaged across simulations within each iteration, and the resulting values were used to update ***C*** according to the following update rule:

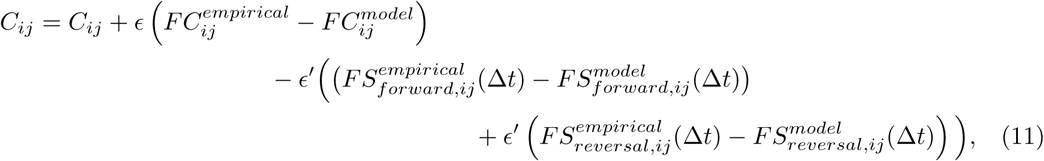

where *ɛ* = 0.0005, *ɛ*^′^ = 0.0001. Additionally, to improve individual model fitting, each subject’s ***C*** was initialized from the group-mean *GEC* matrix of the corresponding menstrual phase. Group-level matrices were first computed using the average empirical matrices for each phase, all initialized from the same ***SC***.

## Statistical analyses

Two separate multilevel linear mixed models (LMMs) were calculated for the hormonal analysis. Estradiol or progesterone served as the dependent variable, while the menstrual cycle phase (early follicular, pre-ovulatory, mid-luteal) was treated as a fixed factor. Participant’s subject identification number was included as a random factor. The statistical model syntax used to examine hormonal level differences across phases was: *hormone level* ∼ *menstrual cycle phase* +(1|*subject*). We adjusted *p*-values using the False Discovery Rate (FDR) [63] method to account for the multiple comparisons. Moreover, we employed LMMs to assess the influence of age, estradiol, and progesterone levels on irreversibility for the whole brain and each resting-state network. Irreversibility values for each node, across participants and cycle phases, served as the dependent variable. Age, progesterone, estradiol, and the interaction between progesterone and estradiol levels were included as fixed effects, while subject ID was treated as a random effect. We determined the model syntax as *Irreversibility value per node* ∼ 1 + *age* + *estradiol* + *progesterone* + *estradiol* ∗ *progesterone* + (1|*subject*). We conducted all multilevel model analyses utilizing R Statistical Software (v4.3.3; [64]) with the package lme4 [65].

### SVM classification for pattern separation

Assessment of group-dependent differences in the individual’s *GEC* matrices was performed using an SVM classifier. This tool circumvents the problem of multiple comparisons, which hampers the assessment of statistically significant differences in high-dimensional matrices such as ours (116 × 116). We evaluated whether the classification accuracy over the 1000 folds was significantly different from chance level with a t-test. The mean classification accuracy across folds was used as a performance metric. Additionally, we extracted the most informative features for each comparison, defined as the ten highest-weighted coefficients in the *GEC*-trained SVM classifier. Each feature represented the effective coupling between an ordered pair of brain regions in the matrix.

## Acknowledgments

E.A. was supported by the FI Grant (no. 2022FI B 00337) funded by the Catalan Agency for Management of University and Research Grants (AGAUR). D.A., B.P. and A.E. were supported by La Fundacío la Marat‘o de TV3 (id 202410-30-31). B.P. and E.H. were supported by the European Research Council (ERC) Starting Grant 850953. G.D. and Y.S.P. were supported by the project NEurological MEchanismS of Injury and Sleep-like cellular dynamics (NEMESIS) (ref. 101071900), funded by the EU ERC Synergy Horizon Europe. G.D. was also supported by the Grant PID2022-136216NB-I00 funded by MICIU/AEI/ 10.13039 / 501100011033 and by”ERDF A way of making Europe”, ERDF, EU. G.D and A.E. were also supported by the project eBRAIN-Health - Actionable Multilevel Health Data (id 101058516), funded by the EU Horizon Europe. A.E. was also supported by the European Union’s Horizon Europe research and innovation programme under the Marie Skl-odowska-Curie Actions (ID: 101207460, NEUROCONTRA, HORIZON-MSCA-2024-PF-01-01), and by the Spanish Ministry of Science, Innovation and Universities with the Jośe Castillejo Grant CAS23/00099. P.D. was supported by the AGAUR FI-SDUR Grant (no. 2022 FISDU 00229).

## Author contributions

Conceptualization: E.A and A.E. Methodology: E.A., Y.S.P, G.D. and A.E. Data analysis: E.A., P.D., M.M., D.A. and A.E. Visualization: E.A., P.D., M.M., and D.A. Data acquisition and preprocessing: E.H. and B.P. Data curation: E.H., B.P. and A.E. Supervision: B.P. and A.E.

Writing—original draft: E.A., D.A, and A.E. Writing—review & editing: all authors.

## Reporting summary

Further information on research design is available in the Nature Research Reporting Summary linked to this article.

## Data availability

Data and scripts are openly available online at http://webapps.ccns.sbg.ac.at/OpenData/ and https://osf.io/23d7x/105(OFS). MR images are available upon reasonable request from the corre-sponding authors.

## Code availability

All code is publicly available in https://github.com/elviradelagua/irreversibility-menstrual-cycle.

